# α-lipoic acid has the potential to normalize copper metabolism, which is dysregulated in Alzheimer’s disease

**DOI:** 10.1101/2021.03.15.435417

**Authors:** Kristel Metsla, Sigrid Kirss, Katrina Laks, Gertrud Sildnik, Mari Palgi, Teele Palumaa, Vello Tõugu, Peep Palumaa

## Abstract

Alzheimer’s disease (AD) is an age-dependent progressive neurodegenerative disorder and the most common cause of dementia. The treatment and prevention of AD present immense yet unmet needs. One of the hallmarks of AD is the formation of extracellular amyloid plaques in the brain, composed of amyloid-beta (Aβ) peptides. Multiple amyloid-targeting drug candidates have recently failed in clinical trials, which creates the necessity to focus also on alternative therapeutic strategies. One factor contributing to the development of AD is dysregulated copper metabolism, reflected in the intracellular copper deficit and excess extracellular copper levels. In the current study, we follow the widely accepted hypothesis that the normalization of copper metabolism leads to the prevention or slowing of the disease and searched for new copper-regulating ligands. We demonstrate that the natural intracellular copper chelator, α-lipoic acid (LA) translocates copper from extracellular to intracellular space in a SH-SY5Y-based neuronal cell model, and is thus suitable to alleviate the intracellular copper deficit characteristic of AD neurons. Furthermore, we show that supplementation with LA protects the *Drosophila melanogaster* model of AD from developing AD phenotype, reflecting in decreased locomotor activity. Collectively, these results provide evidence that LA has the potential to normalize copper metabolism in AD and supports the hypothesis that LA supplementation may serve as a promising cost-effective method for the prevention and/or treatment of AD.

**Significance statement:** Alzheimer’s disease (AD) is a major biomedical concern that requires novel effective prevention and treatment approaches. An early determinant of AD pathology is dysregulated copper metabolism, which initiates the amyloid cascade, induces oxidative stress and impairs the functioning of cellular copper proteins, all contributing to the development of neurodegeneration. We suggest that the natural copper chelator α-lipoic acid (LA) can normalize impaired copper metabolism in AD. We demonstrate that LA promotes the influx of copper into SH-SY5Y cells in a dose-dependent manner. Moreover, we show that LA alleviates the disease phenotype in a Drosophila melanogaster model of AD. Together with previously published data, these results support the hypothesis that LA has the potential for the prevention and treatment of AD.

## Introduction

Alzheimer’s disease (AD) is characterised by the occurrence of amyloid plaques and neurofibrillary tangles in the brain, which results in neurodegeneration and clinical diagnosis of dementia (1). AD is an age-dependent disease affecting approximately 50 million people worldwide and this number is expected to increase dramatically because of the population aging (2). AD is the costliest disease for the developed countries as there is no cure and the patients require long-term support (3-5). It is universally accepted that the prevention and treatment of AD is one of the major current medical problems and the development of effective treatment and prevention strategies will considerably decrease the global healthcare burden (2).

According to the prevalent amyloid cascade hypothesis (6, 7), the formation of amyloid plaques, consisting of amyloid-beta (Aβ) peptides, and the consequent appearance of neurofibrillary tangles, composed of aggregated hyperphosphorylated tau proteins ultimately leads to neurodegeneration in AD. Most therapeutic approaches to AD have focused on targeting Aβ and tau, but unfortunately, none of them have been successful in clinical trials so far (8-10). For this reason, there is the need to target also alternative and more upstream events in AD pathology (11, 12).

There is considerable evidence showing that copper metabolism is dysregulated in AD (13, 14), which may trigger the development of AD. Copper is an essential redox cofactor for more than twenty enzymes with crucial roles in cellular energy production (cytochrome c oxidase (CCO)), antioxidative defence (Cu,Zn-superoxide dismutase-1, (Cu,Zn-SOD-1)), oxidative metabolism (lysyl oxidase, tyrosinase, dopamine β-monooxygenase, peptidylglycine α-amidating monooxygenase etc.) and metabolism of iron (ceruloplasmin (CE)). However, “free” or weakly complexed copper ions generate by interaction with oxygen metabolites reactive oxygen species (ROS), including highly toxic hydroxyl radicals (15). This double-faceted nature of copper ions dictates the requirement for their tight control (16, 17). Dysregulation of copper metabolism, such as deficiency, misdistribution, or excessive accumulation is detrimental and leads to various diseases. (18). Classical examples of excessive accumulation and deficiency of copper are Wilson’s disease (WD) and Menkes disease (MNK), caused by loss-of-function mutations in copper transporters ATP7B (19) and ATP7A (20), respectively. Importantly WD and MD can be treated by correcting the abnormal copper metabolism by using copper chelators (21) or copper supplements (22), accordingly.

Copper metabolism disturbance in AD is characterized by copper misdistribution. AD is accompanied by substantially elevated levels of copper in extracellular space like blood serum and cerebrospinal fluid (CSF) (23-26) and simultaneous copper deficiency in brain tissue (27), which can be detrimental. Similar changes on a smaller scale also occur during the normal aging process (28, 29). According to such a scenario, dysregulation of copper metabolism is an early event in AD pathology and its normalization might have an effective strategy for the prevention and/or treatment of AD. Attempts to regulate copper metabolism in AD, have so far been unsuccessful. According to our opinion, the failures have been caused by several reasons. Generally, so far the attempts have been largely trial and error cases and were not based on a comprehensive understanding of the copper-binding properties of the ligands in comparison with organismal copper proteins and peculiarities of organismal copper metabolism in norm and AD. Secondly - synthetic Cu(II) chelators (30, 31) have been used, which are known to induce undesirable decoppering of the organism (32-34). Third, application of synthetic copper ionophores such as clioquinol leads in contrary to an abnormal increase of cellular copper (35, 36) which becomes toxic (37).

In the current study, we propose molecular tools to normalize copper metabolism in AD based on the systematical knowledge about the metal-binding properties of potential AD drug candidates, which have the potential to normalize distorted copper metabolism in AD. Our lead compound is α-lipoic acid (LA). Earlier work from our group has shown that the reduced dihydro-LA form has a substantial Cu(I)-binding affinity (38). Its affinity for copper is higher than glutathione (GSH) but lower than intracellular copper chaperones and enzymes (Figure 1) (38, 39). LA acts as Cu(I) chelator only in the intracellular space, where it exists in reduced form and may shift the copper equilibrium from extracellular to intracellular. We tested the potential of LA to regulate the cellular copper metabolism in a manner necessary for the treatment of AD. We demonstrate that supplementation with LA promotes the influx of copper into SH-SY5Y cells in a dose-dependent manner. In addition, by using a *Drosophila melanogaster* model of AD, we show that LA can alleviate the disease phenotype of these mutant flies in a negative geotaxis experiment. These results, together with surplus data from the literature support the hypothesis that normalization of copper homeostasis by LA may be a promising avenue for the prevention and/or treatment of AD.

**Figure 1.**
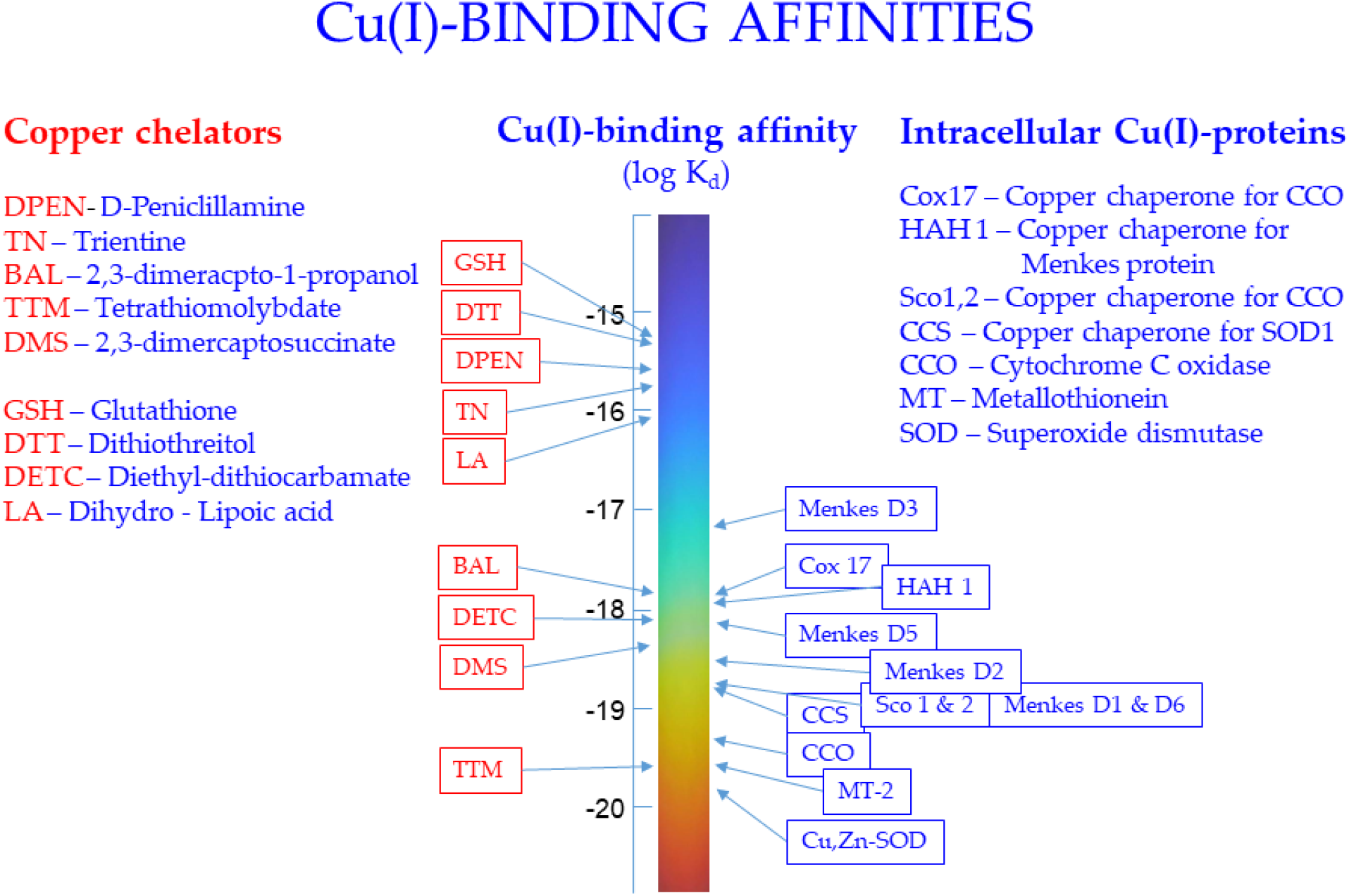
Copper(I)-binding affinities of intracellular Cu(I) proteins and copper chelators according to (39) and (38).

## Results

### The effect of LA on the distribution of copper ions in cell culture

To study whether LA can redistribute copper from extracellular to intracellular environment, we used human neuroblastoma cell line SH-SY5Y, which is a widely used cellular model in neuroscience in the non-differentiated and differentiated form (40). In the current study, we differentiated SH-SY5Y cells with retinoic acid (RA) and brain-derived neurotrophic factor (BDNF), which induces neuronal phenotype of cells reflected in the outgrowth of long neurites and formation of neuronal network (41) (Fig 2, A B). Both types of cells were treated with 5-50 μM LA in the presence of 5 µM CuCl_2_. The results demonstrate that LA promotes the translocation of copper ions from the extracellular environment into cells in a concentration-dependent manner (Fig. 2, C D). The effect was evident already at a low micromolar concentration of LA and was more pronounced in the case of differentiated SH-SY5Y cells (Fig. 2, C D).

**Figure 2.**
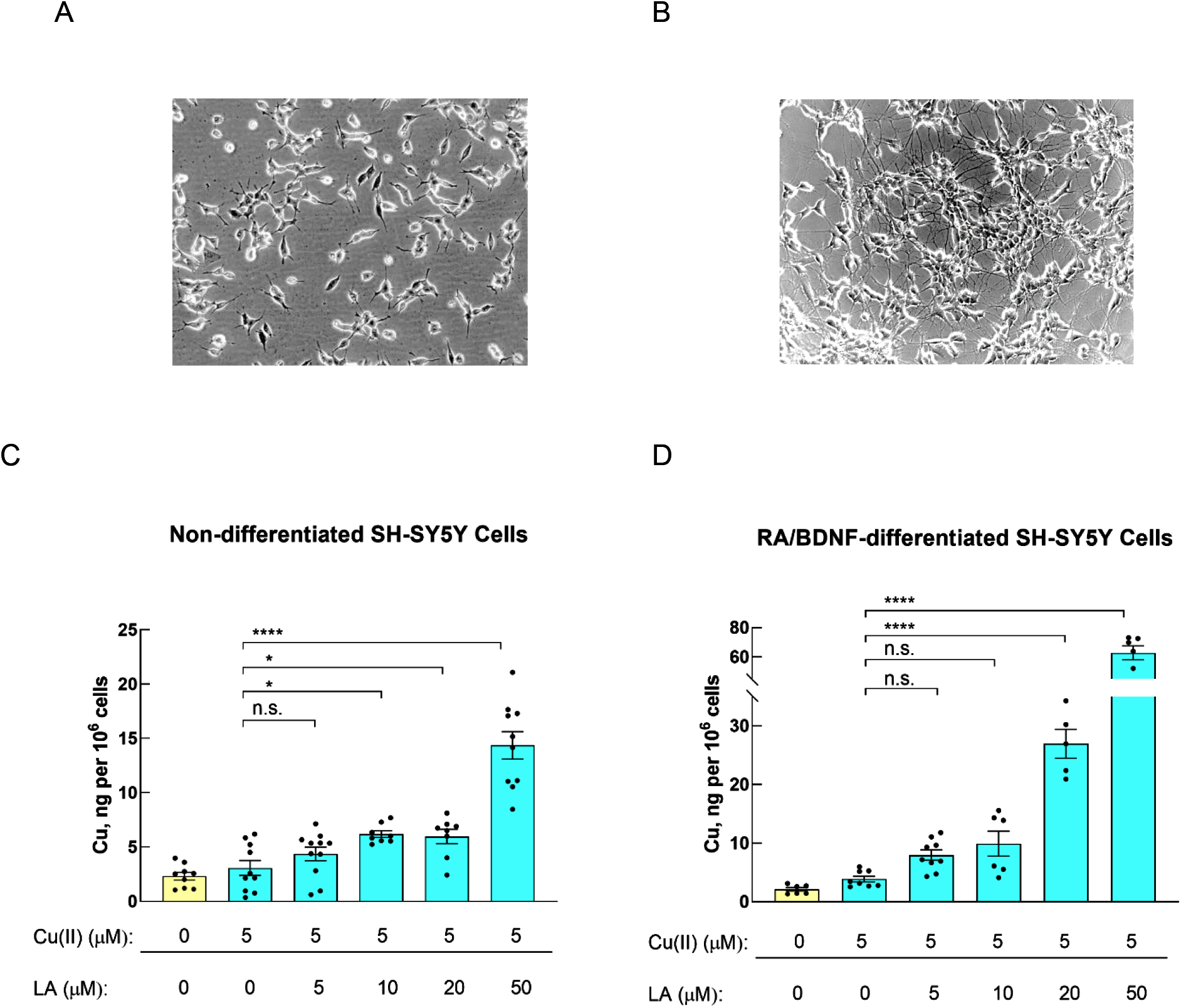
The effect of LA on the cellular copper content in SH-SY5Y cells. Phase contrast images showing the morphology of non-differentiated (A) and differentiated SH-SY5Y cells (B). Non-differentiated (C) and RA/BDNF-differentiated (D) SH-SY5Y cells were treated for 24 h with 0-50 μM LA in the presence of 5 µM CuCl_2_; concentration of Cu, per 10^6^ cells was determined by ICP MS. The columns display the mean ± SEM; n=8-11. One-way ANOVA followed by a Dunnett’s multiple comparisons test at the 0.05 level was used for statistical analysis. Main effect of treatment **** p < 0.0001; * p < 0.05; n.s., not significant.

### The effect of LA on the phenotype of *Drosophila melanogaster* AD model

We used the overexpression of APP or Aβ to model AD in flies (reviewed by (42)). For the ectopic overexpression in *Drosophila*, the two-component Gal4-upstream activating sequence (UAS) system is widely exploited (43). We chose the 30Y-Gal4 driver line with specific expression in the *Drosophila* mushroom body previously shown to be capable to affect also the negative geotaxis (44). For the overexpression we used UAS-APP.Aβ42.D694N.VTR responder line expressing human Aβ with an Iowa mutation from familial AD patients, D23N. The offspring of AD flies and Control flies were provided with standard food (food) and food with LA added (food + LA) after hatching. After 7 days of incubation, the Control flies kept on food and food + LA did not display any difference in the climbing score determined by the negative geotaxis experiment (Fig. 3, B). The climbing score of AD flies incubated on food declined compared to Control flies (Fig. 3, B), whereas AD flies kept on food + LA showed a lower decline, indicating that LA protects AD flies from developing AD phenotype. Analysis of male and female flies separately demonstrated that the effect of LA was statistically significant in both sexes (Fig. 3, C).

**Figure 3.**
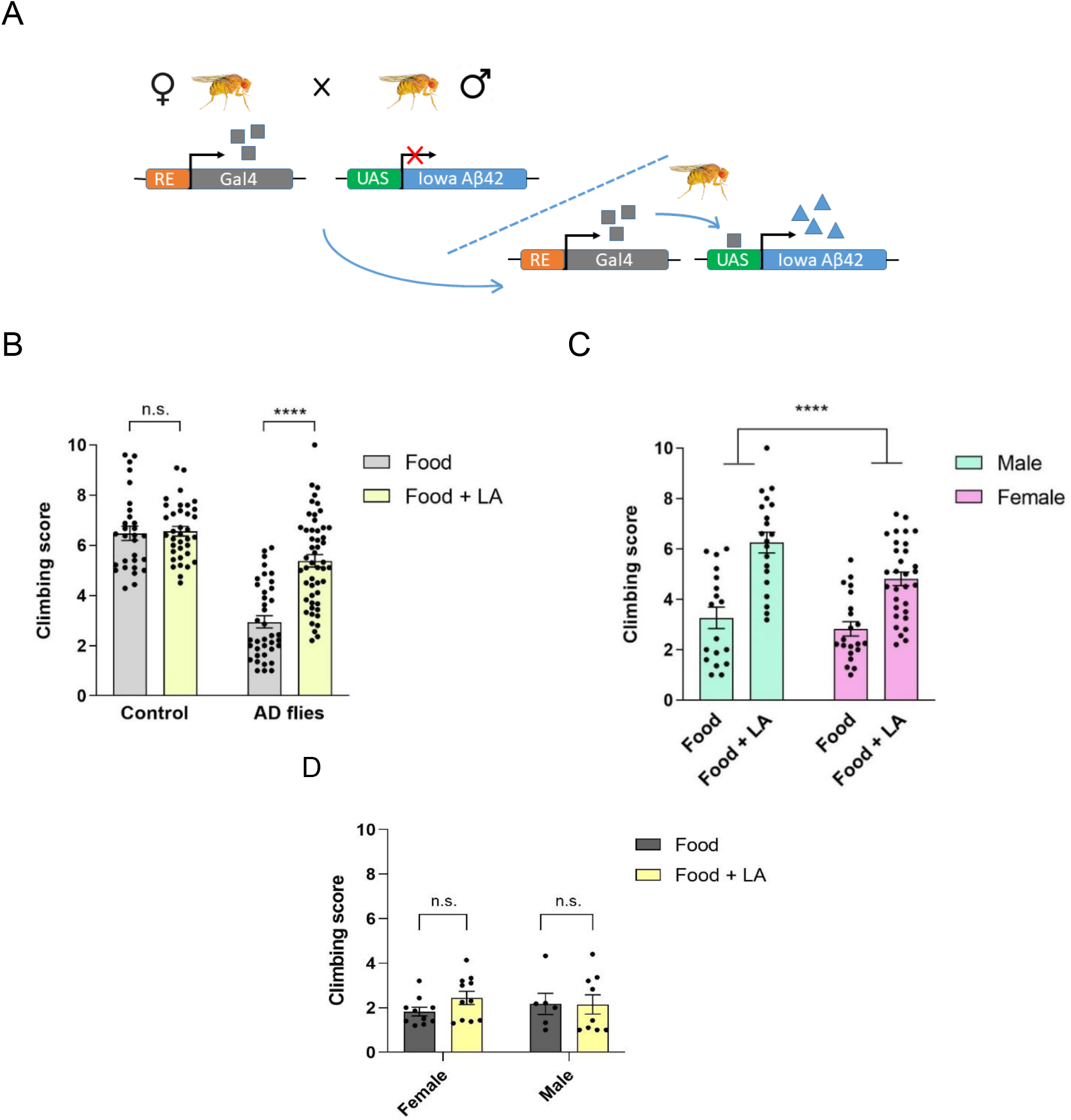
Effect of LA on the climbing ability of Control vs AD flies. **(A)** The crossing of RE (regulatory element 30Y) driven Gal4 flies with flies containing UAS in tandem with Iowa Aβ42 gene (A). AD flies were incubated for 7 days on food or food + LA before a negative geotaxis assay (B, C). AD flies were incubated on food for 7 days and food or food + LA for the next 7 days (D). The climbing score displays the mean ± SEM of climbing distance for n=32-57 in groups, each consisting of 7-10 flies. Two-way ANOVA with Sidak’s multiple comparison correction was used to compare the scores of different experimental groups. Main effect of treatment **** p < 0.0001; n.s., not significant.

**Figure 4.**
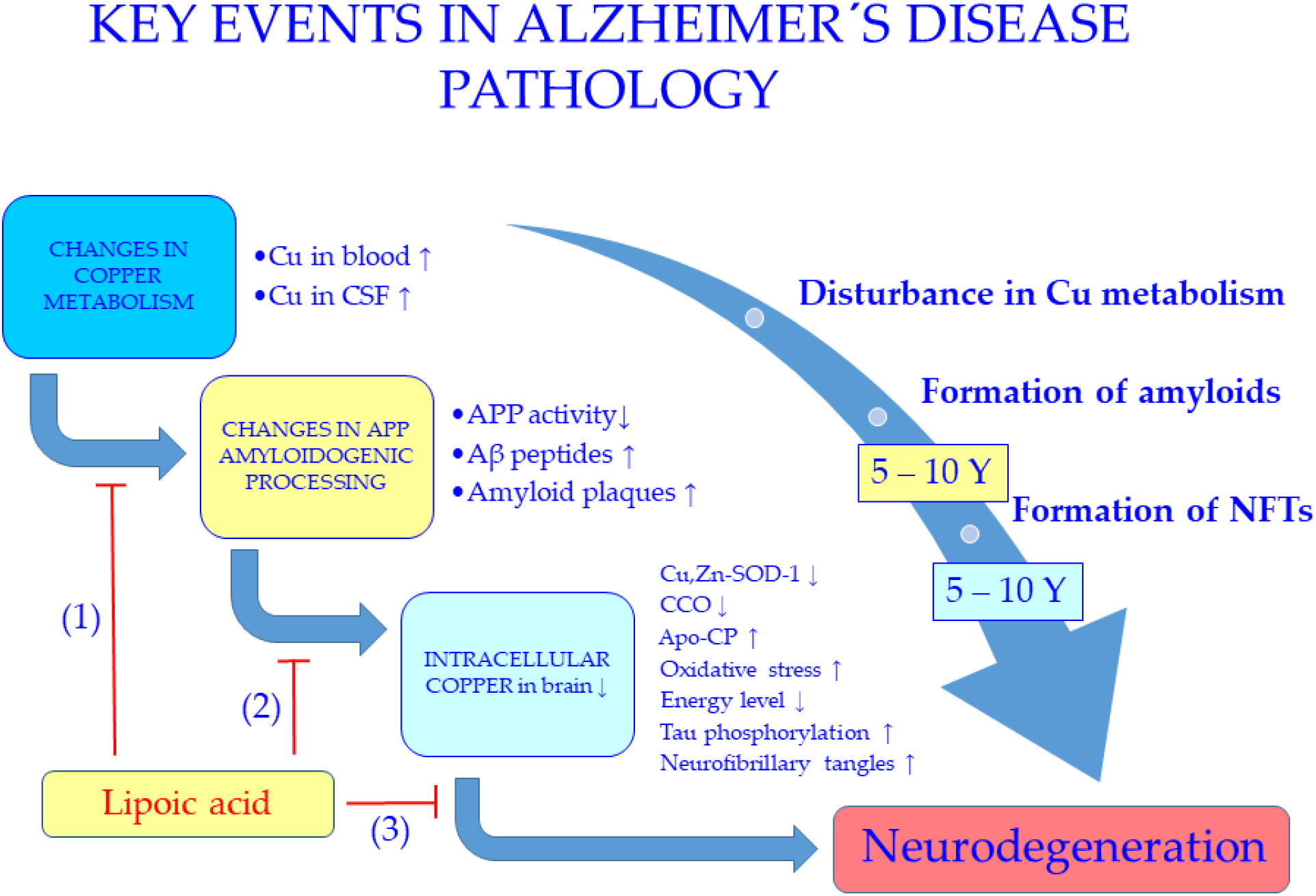
Key events in Alzheimer’s disease pathology. According to our suggestion LA can correct extra- and intracellular copper metabolism (1), suppress amyloid cascade (2) and neurodegeneration (3) characteristic of AD.

We also studied the effect of LA on flies that had already developed the AD phenotype. In this experiment, the AD flies were kept on food for 7 days and half of the flies were thereafter transferred to food + LA for the subsequent 7 days. We found that the supplementation of LA did not affect the results of the negative geotaxis assay in male and female flies (Fig. 3, D) These results suggest that LA is not able to rescue the AD phenotype that has already developed.

## Discussion

In this study, we investigated the molecular tools to normalize copper metabolism, which is dysregulated in AD. Quantitative meta-analyses of numerous independent studies conducted on AD patients have revealed that AD is accompanied by substantially elevated copper levels in serum (23-25) as well as in the CSF (26) and simultaneous copper deficiency in the brain tissue (27). The more precise analysis shows that in the serum and CSF of AD patients mainly non-ceruloplasmin (CP)-bound fraction of copper is increased (45, 46) and in AD brain tissues copper levels are substantially (53 – 70%) decreased in multiple brain regions (47).

The molecular and genetic background of distorted copper metabolism has been also extensively studied in the context of AD. Genetically, AD has been associated with certain variants of *ATP7B*, the copper transporter defective in WD (48). Furthermore, it turned out that copper affects several cellular aspects of AD pathology. First, amyloid precursor protein (APP), a central molecule in AD pathology, the proteolysis of which produces Aβ peptides (49), is a copper-binding protein (50, 51) and its expression (52), oligomeric state (53), cellular localization (54, 55), and proteolysis (56, 57) are copper-regulated. Therefore, the misbalance of copper metabolism may shift the proteolysis of APP towards the amyloidogenic pathway, leading to increased production of Aβ peptides or more amyloidogenic peptide Aβ42, which plays a crucial role in initiating and perpetuating the AD pathologic cascade (7). Second, copper ions bind with relatively high affinity to full-length and truncated Aβ peptides (58-60), accelerating their aggregation (61, 62). Third, copper ions are enriched in Aβ fibrils *in vivo* in AD brains (63) as well as *in vitro* (64) and cause oxidative stress, which is neurotoxic and can cause neurodegeneration (65, 66). Listed evidence indicates that normalising copper metabolism provides an attractive avenue for the treatment and prevention of AD.

Because of the connection between copper metabolism and AD, several attempts have been made to treat AD by modifying copper metabolism. Three different strategies, copper supplementation, chelation, and redistribution have all been tested in laboratory experiments and also in clinical trials. Copper supplementation was attempted with copper orotate (67, 68), chelation with D-penicillamine (34) and copper redistribution with clioquinol (CQ, 5-Chloro-8-hydroxy-7-iodoquinoline) and its derivative PBT2 (5,7-dichloro-8-hydroxy-2-[(dimethylamino)methyl]quinoline) (69, 70). However, copper supplementation showed no effect on the progression of AD phenotype over a 12-month treatment period (67). D-penicillamine promoted decoppering, but did not affect the clinical progression of the disease (34). CQ was also unsuccessful in a clinical trial and has been withdrawn from development due to safety concerns (71, 72). Furthermore, PBT2, although with a favorable safety profile (72), did not reduce amyloid plaques and did not improve the cognitive function of AD patients(73). In addition to clinically tested compounds, numerous other synthetic Cu(II)-binding ligands (30, 31, 74-76), including trientine (triethylene tetramine dihydrochloride) (77), an FDA-approved WD drug, have been proposed for the treatment of AD. Trientine and all other this type of drugs result in a decoppering of the organism, which may further decrease the copper levels in the brain. It is known from MNK that such copper deficiency leads to insufficient metallation of cellular copper proteins and causes neurodegeneration (78). Copper deficiency similar to MNK occurs also in the AD brain and its causative link with AD pathogenesis has been proposed (47). The major clinical neurological feature of MNK is epileptic seizures, which is proposedly linked with deficiency of copper-dependent dopamine β-hydroxylase (79, 80). Studies have shown that patients with AD are also at substantially (6-to 10-fold) increased risk for developing seizures and epilepsy (81). Moreover, epileptiform activity occurs also in AD transgenic mice with neuronal expression of mutant APP and elevated levels of Aβ (82, 83), which supports the hypothesis of the neuropathological role of copper deficiency in AD.

Therefore, copper-binding ligands, which lead to decoppering are not suitable for the normalization of copper dysmetabolism in AD. Rather, substances with the ability to translocate excessive extracellular copper to the intracellular space in the brain is needed. We aimed to normalize copper metabolism by using Cu(I)-binding ligands, which act only in intracellular space and shift the equilibrium of copper distribution from extracellular to intracellular location. Through thorough knowledge of the Cu(I)-binding properties of cellular copper proteins and Cu(I)-binding ligands, we established that dihydro-LA has substantial Cu(I)-binding affinity owing to its two closely located SH groups (38), which can specifically bind Cu(I) ions into the dihedral complex. SH groups of LA are reduced inside the cell and oxidized to a disulfide bond in the extracellular environment thus enabling selective intracellular Cu(I) binding and shifting of equilibrium of copper distribution towards intracellular space.

LA is a natural ligand with its biochemistry and biological effects extensively studied. LA is synthesized enzymatically in the mitochondrion from octanoic acid (84) and is functioning as a cofactor covalently linked to mitochondrial α-ketoacid dehydrogenases (85), essential in mitochondrial energy metabolism. In addition to endogenous synthesis, LA is also absorbed from dietary sources and elicits a unique set of biochemical activities (84). Thus far, the biological effects of LA have been explained mainly by its antioxidant action, however, its potential in detoxification of heavy metals like Hg has also been recognized (86).

LA has also been studied in the context of AD and aging. For example, LA improves the memory of aged nontransgenic (NMRI) mice (87) as well as transgenic AD (Tg2576) mice (88). Moreover, supplementation with LA extends the lifespan of *Drosophila melanogaster* (89) and immunosuppressed mice (90). LA has also clinically proven therapeutic value in the treatment of diabetic polyneuropathy (91). Most importantly, LA has been tested in AD clinical trials. A daily dose of 600 mg showed a positive effect by slowing the progression of cognitive impairment in patients with mild AD (43 patients, trial duration 48 months) (92, 93) and in patients with mild to moderate AD with and without insulin resistance (126 patients, trial duration 16 months) (94). Despite these promising results, clinical trials with LA have not been taken forward because of the largely elusive mechanism of LA action. The therapeutic effect of LA in these trials has mainly been attributed to its antioxidative effect, but its metalloregulatory properties have not been considered or studied.

There are numerous benefits of LA over other synthetic compounds in drug development as well as in further therapeutic use. LA has been approved for the treatment of diabetic polyneuropathy (91) and could be repurposed for therapeutic application in AD (95). Known toxicology and pharmacodynamic profiles of repurposed drugs significantly accelerate the drug development process, decrease the related costs, and increase the probability of success. Finally, LA is a natural compound, approved as a food supplement in many countries and it is freely and cheaply available in pharmacies, which makes it accessible and affordable for AD patients worldwide if its beneficial effects and mechanism of action are approved.

In the current study, we demonstrated that supplementation with LA significantly increases the intracellular copper level of SHSY-5Y cells in a dose-dependent manner. An increase of intracellular copper occurred already at 5 μM concentration of LA and the increase was moderate, which is similar to the decrease of copper level in AD brain tissue (47). The mechanism of LA action in promoting copper influx remains to be established, however, based on available data we suggest that LA acts on copper metabolism through the biochemical mechanism and not through regulation of transcription. Furthermore, by using a *Drosophila melanogaster* transgenic AD model, expressing human Aβ with an Iowa mutation D23N, we showed that early supplementation with LA prevents the development of AD phenotype in transgenic AD flies in a negative geotaxis experiment. Interestingly, LA cannot alleviate already developed phenotype in AD flies. Although the expression of Aβ42 peptide is not directly connected with copper metabolism, there is evidence that the phenotype of these flies is affected by copper. For example, the phenotype of Aβ42 expressing flies is ameliorated through inhibition of high-affinity copper influx transporter Ctr1 orthologues in the fly nervous system (96). In addition, the phenotype of flies expressing Aβ42 specifically in their eyes is changed after mutations in copper transporter ATP7 (97) as well as by adding Cu(II) ions to the food (98). Changes in the phenotype induced by copper supplementation are reversed by copper chelators (98), which shows that copper metabolism is distorted in the AD fly model expressing Aβ42 peptides and its normalization has a beneficial effect on the phenotype of flies. Therefore, the beneficial effect of LA on AD flies may also be connected with the normalization of copper metabolism, however, it has to be confirmed in follow-up studies.

In conclusion, we propose a mechanism of LA action in AD (Fig. 5), which states that LA can normalize copper distribution between extra and intracellular locations, which inhibits amyloidogenic processing of APP, decoppering of cellular copper proteins and other downstream processes including neurodegeneration. The detailed molecular events remain to be determined, however, our results and the data on the beneficial effects of LA in AD support its applicability primarily for the prevention and inhibiting the development of AD pathology, which is desperately needed. It is estimated that the introduction in 2025 of an agent that delays AD onset by 5 years would decrease the total number of patients with AD by 50% in 2050 (99). LA may prove to be an effective strategy to prevent the development of AD and slow its progression and double-blind clinical experiments are warranted.

## Materials and methods

### Cell culture experiments

Human SH-SY5Y neuroblastoma cells (ATCC) were grown in Dulbecco’s Modified Eagle Medium (DMEM) (Gibco) supplemented with 10% fetal bovine serum (FBS) (Gibco) and 1% penicillin and streptomycin (PEST) (Gibco) at 37 °C in a humidified atmosphere containing 5% CO_2_. The culture medium was changed every 2 to 3 days and the cells were split every 5 to 7 days using 0.25% Trypsin-EDTA (Gibco), up to 20 times. Cells were plated with a density of 2 × 10^5^ cells/ml into white clear-bottom 6-well plates (Greiner Bio-One) 2 ml per well for inductively coupled plasma mass spectrometry (ICP-MS) experiments. SH-SY5Y cells were differentiated in cell culture plates, using the following differentiation protocol: cells were pre - differentiated with 10 μM RA in full medium for 4 days, followed by differentiation with 50 ng/mL BDNF (Alomone Labs) in serum-free medium for 2 days (100). Phase-contrast images of cells were taken using Zeiss Axiovert 200M microscope with 20x objective. For cell count measurement 10 μl of cell suspension was mixed with an equal volume of trypan blue stain, pipetted into Countless cell counting chamber slide, and inserted into the Countess Automated Cell Counter (Invitrogen). Non-differentiated and differentiated SH-SY5Y cells were treated for 24 h with 5 μM CuCl_2_ (Sigma-Aldrich) in the presence of 5 – 50 μM LA (Sigma Aldrich).

### ICP-MS analysis

Differentiated SH-SY5Y cells were collected from 6-well plate wells into acid-washed 15 ml tubes and centrifuged for 1 min at 10 000 xg to separate the medium from cells. 100 μl of supernatant was collected into acid-washed Eppendorf tubes to use as a medium sample. Cells were washed twice with 500 μl PBS (Sigma Aldrich), and a sample of 10 μl from cell suspension was taken to measure cell count with Countess Automated Cell Counter as previously described. Cells in PBS were centrifuged for 3 min at 10 000 xg to separate cells from PBS. PBS was discarded and the obtained cell samples and collected medium samples were stored at-20 °C until the ICP-MS analysis.

One day before ICP-MS analysis, 100 μl of 68% HNO_3_ was added to 100 μl of collected cell culture medium and separated cells. Acidified samples were incubated for 24 h at room temperature. For ICP-MS analysis samples were diluted to 3.4% HNO_3_ [71, 72]. Ultrapure Milli-Q water with a resistivity of 18.2 MΩ/cm, produced by a Merck Millipore Direct-Q & Direct–Q UV water purification system (Merck KGaA, Darmstadt, Germany), was used for all sample preparations.

ICP-MS analyses for Cu-63 were performed on an Agilent 7800 series ICP-MS instrument (Agilent Technologies, Santa Clara, USA) and Agilent SPS-4 autosampler was used for sample introduction. For instrument control and data acquisition, ICP-MS MassHunter 4.4 software Version C.01.04 from Agilent was used. ICP-MS analysis was performed in peak-hopping mode, 6 points per peak, 100 scans per replicate, 3 replicates per sample, and the instrument was operated under general matrix working mode under the following conditions: RF power 1550 W, nebulizer gas flow 1.05 l/min, the plasma gas flow 15 l/min, nebulizer type: MicroMist. Elements monitored: Sc-45 (internal standard) and Cu-63. The ICP-MS apparatus was calibrated using multielement calibration standard 2A in 2% HNO_3_ (Agilent Technologies, USA) containing Ag, Al, As, Ba, Be, Ca, Cd, Co, Cr, Cs, Cu, Fe, Ga, K, Li, Mg, Mn, Na, Ni, Pb, Rb, Se, Sr, Tl, U, V, Zn at levels 0, 0.1, 0.5, 1, 5, 10, 25 and 50 ppb with Sc-45 (ICP-MS internal standard mix 1 ug/mL in 2% HNO_3_, Agilent Technologies) as the internal standard for Cu-63.

### *Drosophila melanogaster* experiments

#### Used stocks, maintenance and husbandry of *Drosophila melanogaster*

The following stocks were used for the experiments: 30Y-Gal4 driver line (a gift from Mark Fortini) (101) and UAS-APP.Aβ42.D694N.VTR (Iowa) responder line (Singh, C., Mahoney, M. (2011), Bloomington Drosophila Stock Center, BDSC_33779). Fly stocks were maintained at 25 °C (12h:12h L:D cycle; 60–70% RH) on malt-semolina based food prepared from 6,5 g agar, 38 g semolina, 70,5 g malt flour, 17,5 g dry yeast, 5,9 ml nipagin (Tegosept 30%; 30g/100ml 94% EtOH; Dutcher Scientific), and 6,8 ml propionic acid (Sigma) per 1000 ml water. *Drosophila* crosses were performed at 29 °C to get a higher activity of *Gal4-UAS* system. By crossing the two lines, the offspring (AD flies) have Aβ42 D23N overexpression driven by 30Y-Gal4 pattern in the fruit fly brain. For controls we used the pure UAS-APP.Aβ42.D694N.VTR line inactive in the absence of Gal4.

### LA feeding regimen

To study the effect of LA on the locomotor activity of AD flies, dietary supplementation with LA was performed. Briefly, LA was dissolved in 96% EtOH for a 200 mM stock solution and added into the previously described food at a final concentration of 2 mM (food + LA). Control food (food) contained the same amount of EtOH that was used to produce LA-supplemented food. Adult male and female flies were separated and added to LA-food and food within 24 h after eclosion and aged at 29 °C for 7 days before the experiments. To study the effect of LA on the developed AD phenotype, separated male and female flies were maintained on regular food for 4 or 7 days and transferred to food + LA and food for further 7 days. In all experiments, flies were transferred to fresh food after every 2 days.

### Negative geotaxis assay for *Drosophila melanogaster*

Before the test, 7-10 flies of each group were transferred by tapping into empty 15 ml vials, covered with another upside-down vial, and vials were connected with transparent tape. Obtained vials were placed in front of a 20 cm high background that was divided into 10 equal spaces. For the measurement, flies were knocked to the bottom of the vial three times, and photos were taken after 10 seconds. The climbing height of each fly was registered, and an average climbing distance score was calculated.

### Statistical analyses

Statistical analyses were performed using GraphPad Prism 8. Data of ICP-MS was analyzed using a one-way analysis of variance (ANOVA) with the posthoc Dunnett’s multiple comparison test. The figures display the mean ± standard error of the mean (SEM). Negative geotaxis scores were averaged per vial (5-20 flies per vial) and the mean score of each vial was treated as an individual replicate for further analysis. Two-way ANOVA with Sidak’s multiple comparison correction was used to compare the scores of different experimental groups. Data on graphs are presented as mean ± SEM and a *p* value of ≤0.05 was considered as statistically significant. Statistical significance of *p* ≤ 0.05 is represented as *, *p* ≤ 0.01 as **, *p* ≤ 0.001 as ***, and *p* ≤ 0.0001 as ****.

## Abbreviations

Aβ: amyloid-beta peptide
AD: Alzheimer’s disease
APP: amyloid precursor protein
BDNF: brain-derived neurotrophic factor
CCO: cytochrome c oxidase
CE: ceruloplasmin
CQ: clioquinol, 5-Chloro-8-hydroxy-7-iodoquinoline
CSF: cerebrospinal fluid
LA: α-lipoic acid
MNK: Menkes disease
PBT2: 5,7-dichloro-8-hydroxy-2-[(dimethylamino)methyl]quinoline
RA: retinoic acid
SOD: superoxide dismutase
UAS: upstream activating sequence
WD: Wilson disease

## Acknowledgments

This work was supported by the Tallinn University of Technology and by the Estonian Research Council grant to PP (PRG 1289).

## Author Contributions

Idea, P.P., Conceptualization, P.P., V.T., T.P., and M.P.; Methodology, P.P., K.M., S. K., M. P., K.L., and V.T.; Investigation, K.M., S.K., K.L., and G.S.; Formal analysis, K.M., S.K., K.L., G.S. and T.P.; Writing – Original Draft, P.P.; Writing – Review & Editing, all Authors; Funding Acquisition, P.P.; Resources, P.P. and M.P.

